# Mechanisms of allosteric and mixed mode aromatase inhibitors

**DOI:** 10.1101/2020.10.15.340745

**Authors:** Samson A. Souza, Abby Held, Wenjie Lu, Brendan Drouhard, Bryant Avila, Raul Leyva-Montes, Michelle Hu, Bill R. Miller, Ho Leung Ng

**Affiliations:** Department of Biochemistry and Molecular Biophysics, Kansas State University, Manhattan, KS; Department of Chemistry, Truman State University, Kirksville, MO; Department of Chemistry, University of Hawai‘i at Mānoa, Honolulu, HI

**Keywords:** Aromatase, aromatase inhibitor, P450 inhibitor, allosteric inhibitor

## Abstract

Aromatase (Cyp19) catalyzes the last biosynthetic step of estrogens in mammals and is a primary drug target for hormone-related breast cancer. However, treatment with aromatase inhibitors is often associated with adverse effects and drug resistance. In this study, we used virtual screening targeting a predicted cytochrome P450 reductase binding site on aromatase to discover four novel non-steroidal aromatase inhibitors. The inhibitors have potencies comparable to the noncompetitive tamoxifen metabolite, endoxifen. Our two most potent inhibitors, AR11 and AR13, exhibit both mixed-type and competitive-type inhibition. The cytochrome P450 reductase-Cyp19 coupling interface likely acts as a transient binding site. Our modeling shows that our inhibitors bind better at different sites near the catalytic site. Our results predict the location of multiple ligand binding sites on aromatase. The combination of modeling and experimental results supports the important role of the reductase binding interface as a low affinity, promiscuous ligand binding site. Our new inhibitors may be useful as alternative chemical scaffolds that may show different adverse effects profiles than current clinically used aromatase inhibitors.

## Introduction

Aromatase (Cyp19) catalyzes the three-step transformation of androgens to estrogens in mammals by a mechanism that is well-characterized. The first two steps produce a C-19 gem-diol ^1^. 1β-hydrogen abstraction and electron transfer furnish the A-ring aromatization and loss of a formate molecule to complete a single turnover ^2^. As such, aromatase inhibition is a common strategy to treat patients with hormone-dependent breast cancer ^3^. Third-generation aromatase inhibitors (AIs) are used as primary therapeutics and long-term adjuvants in postmenopausal women with breast cancer.

AIs in clinical use include the non-steroidal AIs (NSAIs), anastrozole and letrozole, and the steroidal AI, exemestane, which interacts covalently at the active site ^4,5^. The most potent NSAI, letrozole, binds a distinct site exhibiting both noncompetitive and mixed-mode type Michaelis inhibition ^6^. This type of behavior is also observed for the noncompetitive AI endoxifen (K_i_ = 4 μM), a potent metabolite of tamoxifen, a clinically-used estrogen receptor α (ERα) antagonist ^7^. Metabolic N-demethylation of endoxifen produces a competitive AI, norendoxifen (K_i_ = 35 nM) ^8^. This demonstrates two important points. First, small chemical modifications of inhibitors (letrozole to anastrozole, and endoxifen to norendoxifen) can change the inhibitory mechanism in unexpected ways. Second, Cyp19 can be allosterically modulated with high potency. The mechanism of mixed mode inhibition for Cyp19 and other cytochrome P450 (CYP450) enzymes is unclear.

Steroidal AIs interact at the active site, also known as the distal heme site, with high efficacy. However, steroidal analogs frequently exert similar adverse effects. Cyp19 inhibition at an alternative allosteric site provides opportunities for the discovery of novel NSAIs chemically distinct from current therapeutics and with different toxicity profiles.

In this work, we investigate the binding of small molecule inhibitors to the Cyp19 heme distal site, the heme proximal site (the predicted site of CPR binding), and the substrate access channel through a combination of experimental and computational approaches. We first predict and model the interfacial contacts of cytochrome P450 reductase (CPR) with Cyp19. We performed virtual screening against a library of over a million compounds to identify potential inhibitors. We use molecular dynamics simulations to model how they interact at the CPR-Cyp19 interface. We then provide experimental enzyme inhibition data for four new NSAIs we discovered, AR11, AR13, AR19, and AR20. In addition, we use optical absorption spectra to characterize the effects of inhibition on the heme chemical environment. We characterize the inhibition modes of AR11, AR13, AR19, and AR20, and their likeliest binding sites closer to the catalytic site. The cytochrome P450 reductase binding site is likely a transient, low-affinity binding site for multiple ligands.

## Materials and Methods

### Protein-protein docking

For protein-protein docking, we used the crystal structures of human CYP19 (PDB 4KQ8) with a human-yeast chimeric enzyme ^9^ (PDB 3FJO) in an open conformation. There is currently no full-length human CPR crystal structure with an exposed FMN-containing face. Human CPR has an intrinsically dynamic hinge region causing the N-and C-terminal domains to adopt a similar open conformation ^10,11^. The chimeric protein maintains functionality reducing both cytochrome c and human P450s ^9^. The Haddock^12^ webserver was used for the interface prediction-driven docking ^13^ of Cyp19 (PDB 4KQ8) to CPR. Haddock uses the CPORT ^13–18^ ensemble with multiple interface predictors to predict the interfacial residues to input as restraints to return high scoring clusters of complexes. Residues that are solvent-exposed and predicted to reside at the CYP19-CPR interface were defined as “Active” for Haddock docking, and surrounding solvent-exposed residues were defined as “Passive” (Table S1). The Gibbs free energy and dissociation constant estimates were calculated by the PRODIGY prediction webserver ^19,20^. Conserved Cyp19 surfaces were calculated using Bayes’ theorem with the ConSurf server ^21,22^. Human Cyp19 (PDB 4KQ8) was queried against 150 homologs across different P450 families by setting a 30 % maximum and 10 % minimum sequence identity cutoff.

### Virtual screening and docking

Idock ^23,24^ was used to virtually screen a library of over one million drug-like molecules from the ZINC database ^25^. The highest scoring molecules were rescored with the DSX knowledge-based scoring function, which has been shown to be more predictive of binding affinity than docking scoring functions ^26^. Compounds were assessed by the interactions made at the proximal heme site and their DSX scores. The top hits were selected for *in vitro* enzyme inhibition screening. Compounds that exhibited moderate to potent anti-aromatase activity (except AR13, for which we had identified the binding site by optical absorption assays) were docked against a crystal structure of Cyp19 (PDB 3S79) with a more stringent protocol for simulations, involving 10 independent docking runs with Autodock Vina using different random seeds ^27^. Autodock Vina exhaustiveness was set to 128, and the number of binding modes was capped at 20 per run. Visual inspection by VMD ^28^ of clustered docking results was used to predict the best binding mode to be used for MD simulations. The stereochemistry of AR13 was ambiguous from the vendor’s (Enamine) molecular description. Both the cis- and trans-cyclopropane forms were used for docking and further modeling.

### Molecular dynamics simulations

The AMBER16 ^29^ molecular dynamics package was used to produce MD simulations. Parameters for the unliganded heme described by Shahrock et al. were used ^30^. The FF14SB ^31^ and GAFF ^32^ forcefields were used for protein and ligands, respectively. Simulations were carried out with the TIP3P explicit solvent model ^33^ in an octahedral box truncated 11 Å from the protein surface. Chloride ions were added for charge neutralization of the system. Energy minimizations were achieved with an 8 Å cutoff for the non-bonded energy term for each atom. A subsequent 2 ns heat step to 310 K with the SHAKE algorithm fixed bond lengths involving hydrogen atoms. A 3.5 ns equilibration step preceded the 1 μs production runs. Simulations were run in triplicate under constant temperature (310 K, Langevin thermostat) and pressure (1 atm, Berendsen barostat) conditions using periodic boundaries. Trajectories were processed using the AMBER cpptraj tools ^34^. The binding free energies and decomposition free energies were calculated using MMPBSA.py ^35^.

### Molecular graphics depictions and regression analyses

Standard molecular graphics were visualized with Pymol (The PyMOL Molecular Graphics System, Version 2.0 Schrödinger, LLC.) Conservation maps were generated with UCSF Chimera ^36^ software. All regression analyses were performed in GraphPad Prism version 8.0.0 for Windows, GraphPad Software, San Diego, California USA. GraphPad functions used for curve-fitting are reported in the appropriate methods sections.

### IC50 assays of top candidates

Compounds were purchased from vendors including Enamine and ChemBridge. We resolved the stereochemistry of compound AR13 using 1D and 2D NMR, which supported a trans configuration at the cyclopropane moiety. In brief, very weak through-space coupling and a ^3^J value of 3.4 supported protons with an uneclipsed dihedral (Figure S1, S2).

Enzyme activity was measured by monitoring the conversion of exogenous substrate 7-methoxy-4-(trifluoromethyl)coumarin (MFC) to its fluorescent product 7-hydroxy-4-(trifluoromethyl)coumarin (HFC). A Cyp19/MFC high-throughput screening kit (Corning) was used to measure inhibition of aromatase activity for compounds AR11 and AR13. Briefly, 2X NADPH regeneration system (16.25 μM NADP+, 825 μM MgCl_2_, 825 μM glucose-6-phosphate, 0.4 U/mL G6P dehydrogenase) was prewarmed with inhibitors (1:2 serial dilutions) at 37°C for 10 minutes in black 96-well plates. Reactions were initiated with prewarmed 2X enzyme-substrate mix (15 nM P450 microsomes enriched with oxidoreductase, 50 μM MFC) and incubated at 37°C for 30 minutes. The 200 μL reaction mixtures were terminated with 75 μL 0.5 M Tris-base (80% ACN). A FluoDia T70 plate reader measured HFC product formation with excitation/emission filters of 400/530 nm. Reactions were performed in duplicate then repeated twice more on separate days.

The reaction conditions were repeated for measuring the inhibitory activity of compounds AR19 and AR20. Modifications are highlighted here. Supersomes containing Cyp19 + CPR (Corning), NADPH regeneration system (Corning), MFC (Chemodex), and ketoconazole (Selleck Chemicals) were used for the reaction mixture. Temperature-controlled incubations of the 96-well plate were carried out with a dry-plate. A Tecan fluorescent plate reader measured the fluorescent product in the circle-read mode at the optimum gain with excitation/emission filters of 405/535 nm. Reactions were measured in duplicate.

Reaction blanks were used for data corrections and these results were normalized to the fluorescence response in the absence of inhibitor. Analyses were performed in GraphPad Prism 8 software, and data were fit to a 4- parameter logistic model on semilog axes.

### Soret shifts by absorption spectroscopy

Codon-optimized Cyp19A1 cDNA in the pCW expression vector was a generous gift from the F. Peter Guengerich lab (Vanderbilt University). Preparation of the DNA construct is detailed by Sohl and Guengerich ^1^. Recombinant protein was produced, purified, and characterized by methods detailed in the supplementary materials section.

Purified Cyp19 was diluted with 100 mM potassium phosphate buffer (pH 7.4) to 50 μL to a final concentration of 2–3 μM P450. Inhibitors were titrated such that the endpoint would not exceed 3 % ACN. Absorption scans were read with a single-beam Agilent 8453 UV-Vis spectrophotometer after resuspension and a 10-minute incubation period at 25 °C. Buffer A components, endoxifen, AR11, AR13, AR19, and AR20 did not contribute to hyperchromic shifts in the Soret peak region.

### AR11 and AR13 kinetics assay

Cyp19 + reductase microsomal preparations (BTI-TN-5B1-4, Corning) were used to measure the conversion of substrate MFC to HFC. Reactions contained 96 μL of NADPH regeneration solution and 4 μL of inhibitor from serial stocks. Mixtures were prewarmed at 37°C in black 96-well round-bottom plates. Reactions were initiated with a single stream of 100 μL prewarmed enzyme-substrate solution in sequence. Parafilm and aluminum foil were applied to plates before incubation within the linear range at 37°C for 20 minutes. The reactions were quenched with 75 μL of 0.5 M Tris-base (80% ACN). HFC was measured with a Tecan plate reader in circle-read mode at 52 gain with excitation/emission filters of 405/535 nm. Data was generated from 10 reads and a 40 μs integration time. Reaction mixtures contained 100 mM PPB (pH 7.4), 10 nM P450, 0.325 mM NADP^+^, 0.825 mM glucose 6-phosphate, 0.825 MgCl_2_, 0.1 U/mL glucose 6-phosphate dehydrogenase, MFC (9.9, 14.8, 22.2, 33.3, and 50 μM), and various concentrations of inhibitor. Ligand sequestration was minimized by working at the lowest enzyme/microsomal concentration that would return a quantifiable fluorescence response. At steady-state conditions, the effect on MFC is negligible since the MFC concentration is greater than 1000-fold that of the enzyme. We used an NADPH regeneration system to minimize the effects of NADPH depletion on the fluorescence response. GraphPad Prism 8 software was used for generating regression curves and analyzing data. 5 × 5 Lineweaver-Burk and 5 × 4 Dixon plots were used to diagnose AR11 and AR13 inhibitory modes. Nonlinear regression curves were fit to mixed- or competitive-type Michaelis functions to return kinetic constants.

## Results

### Predicted Cyp19-CPR interactions

P450s share a conserved fold with low sequence identity between different enzymes. Nearly all the conserved residues are in the buried regions. We used ConSurf, which identifies conserved surfaces ^21^, to find four major surface-exposed sites that are conserved across different P450 families (Figure 1).

**Figure 1.**
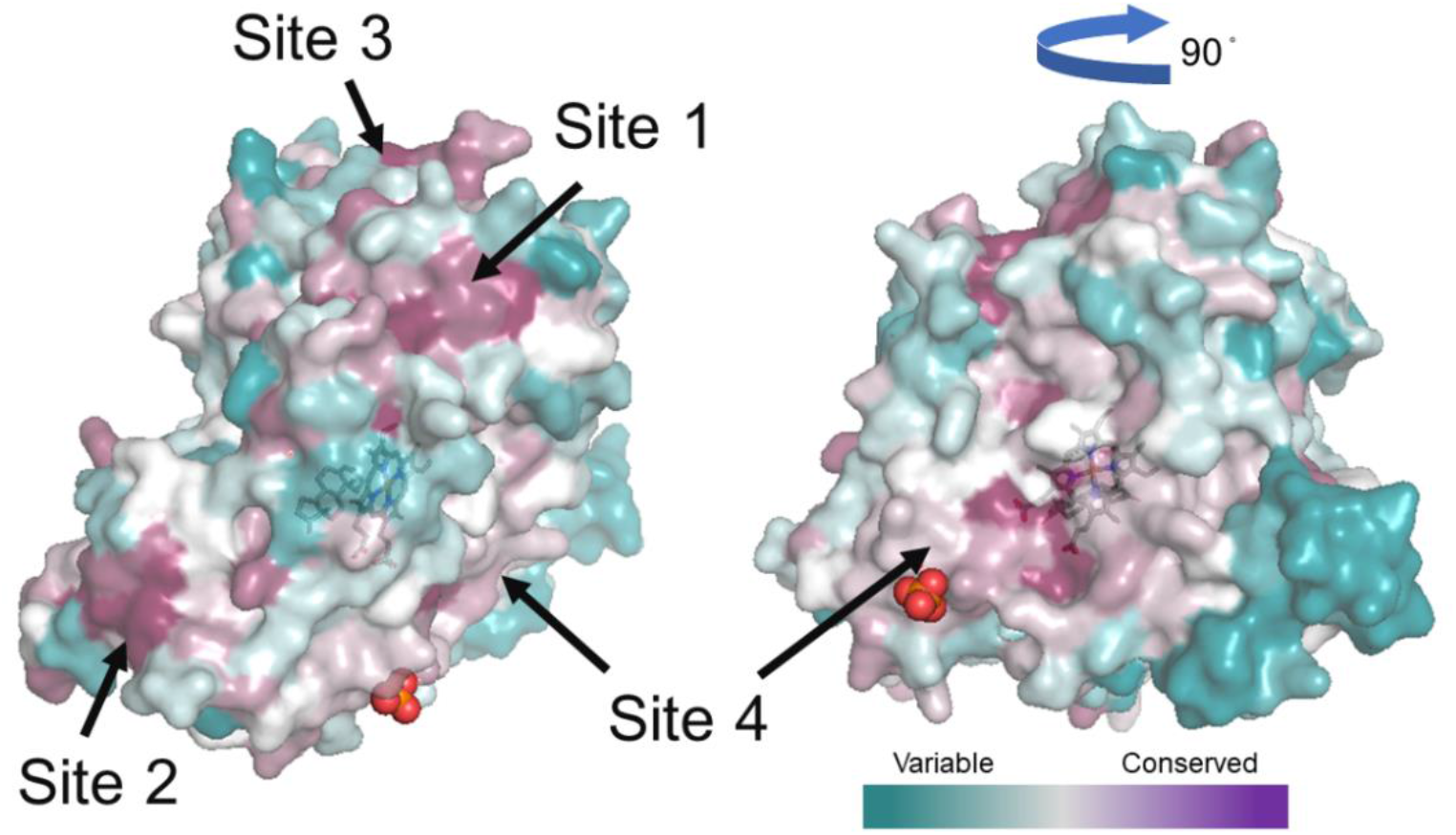
Surface representation of variable and conserved regions of aromatase coded as a cyan to magenta color-gradient. The four conserved regions are labeled as Site 1 – 4. Site 2 is the N-terminus linked to the transmembrane segment. Site 4 corresponds to the proximal heme site.

Only two of the sites were predicted by the protein-protein interface algorithm, CPORT ^13^, to actively participate in protein-protein interactions (sites 2 and 4 in Figure 1). They correspond to the N-terminal region (αA’ and β1- 2) and the proximal heme site. The likeliest CPR binding surface was selected based on three criteria. Firstly, the proximity of the N5 atom of FMN to the iron center of the heme group should be within a distance that is physiologically sound. Secondly, the orientation of the N-terminus of the reductase and Cyp19 should be positioned in the same direction since they are truncations of transmembrane segments. Lastly, only the highest-scoring clusters from docking aromatase with CPR with Haddock ^12^ were assessed. Only site 4, corresponding to the proximal heme region, fulfilled all three criteria.

The probable binding mode contrasts with that of the crystal structure of *B. megaterium* P450BM3 fusion (PDB 1BVY). The N5 to iron distance in our structure is 14.4 Å, whereas the distance is 22.7 Å in P450BM3. This is within the 14 - 15 Å threshold limit for electron transfer in most physiological processes ^37^. The possibility of through-bond tunneling at much longer distances in P450 BM3 was previously refuted due to faster experimental kinetic rates than predicted from theoretical models ^38^. Physicochemical descriptions of the modeled and fusion P450-CPR complexes are compared in Table S2. In docking Cyp19 against CPR in a closed conformation (PDB 3QE2), the closest N5 to iron distance was 34.9 Å. The negative electrostatic potential of the FMN domain interacts with the positive potential of the FAD/NADPH domains. In this conformation, FADH_2_ is in a closer proximity to reduce the FMN cofactor. In the proposed end-on interaction, 3FJO adopts an extended conformation to expose the buried FMN-binding interface for the reduction of Cyp19 at the proximal heme site (Figure 2, S3).

**Figure 2.**
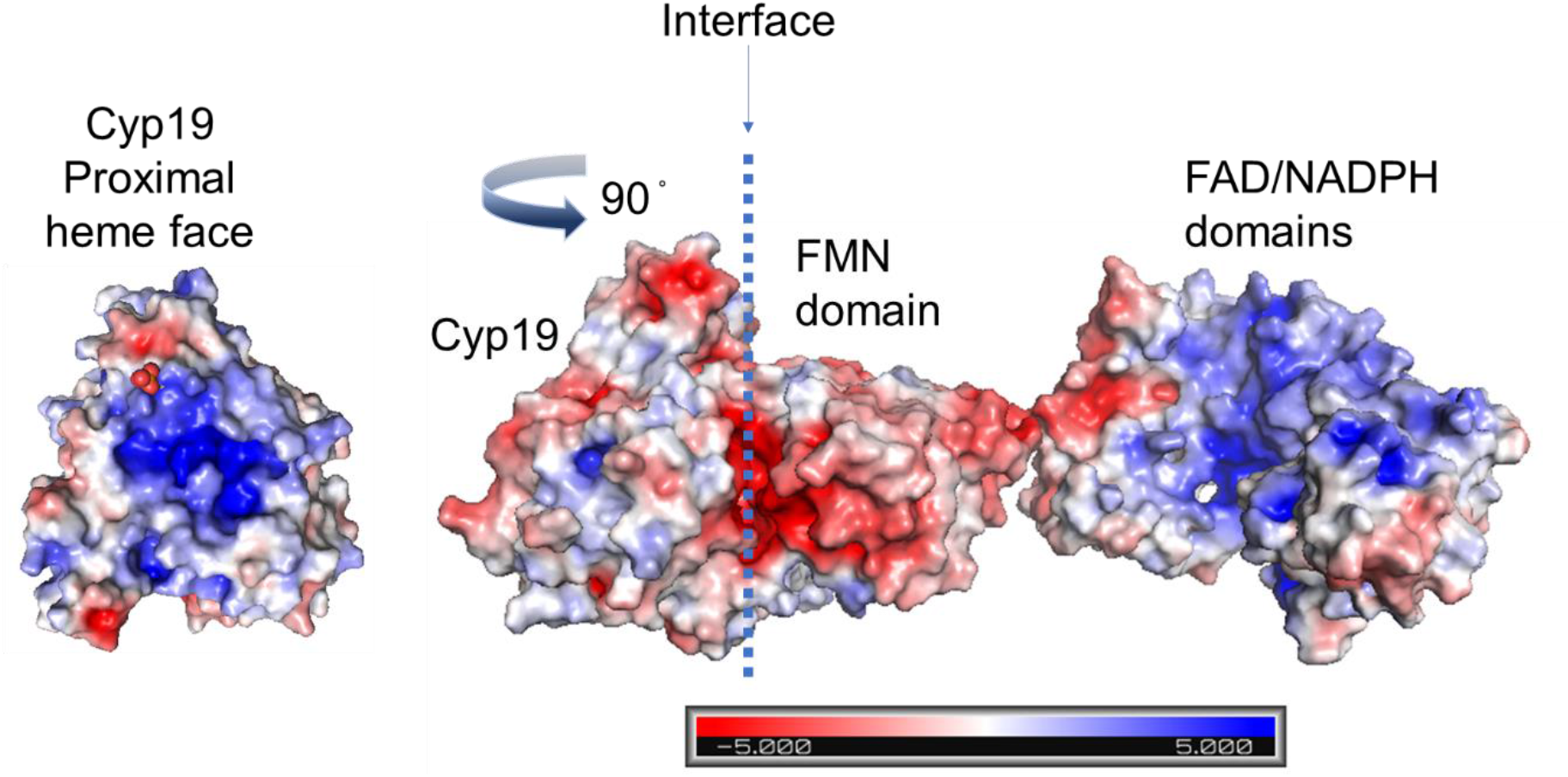
Electrostatic potential surface map of the proposed end-on binding mode of Cyp19 (PDB 4KQ8) in complex with CPR (PDB 3FJO) in an open conformation. Negatively charged potentials (red) to positive potentials (blue) are represented as a color gradient with neutral (gray) regions.

At the Cyp19-CPR interface there are 17 total polar-polar, polar-charged, and charged-charged bond pairs. The most important of these contacts involve sidechains from the Cyp19 residues K108, Y424, K440, and Y441 (Figure 3, Table S2).

**Figure 3.**
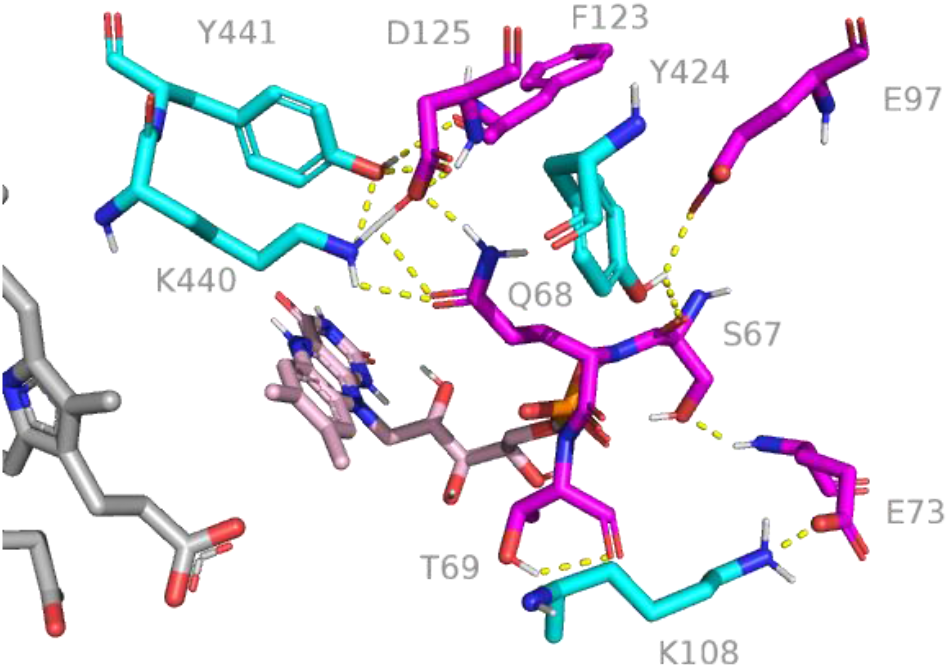
Predicted electrostatic contacts (yellow dotted trace) between Cyp19 (cyan) and the CPR FMN domain (magenta). The FMN cofactor (pink) and heme (gray) are within the 15 Å limit of electron tunneling processes.

### IC50 values of top hits

The DSX (knowledge-based scoring function)^26^ scores and chemical structures of the top four hit compounds are provided in Figure 4 and Table S3. These compounds showed IC_50_ values < 70 μM. AR13 showed 3-fold increased potency over the control inhibitor, ketoconazole, an antifungal with IC_50_ = 3.08 μM. The IC_50_ value we report for AR19, 66 μM, was fit to a Hill coefficient of 1 with a predicted 14% activity at saturation. The R^2^ is suboptimal due to the poor solubility of AR19. AR11 and AR20 exhibited potency comparable to an active tamoxifen metabolite, endoxifen. Dose-response curves and activity data are presented in Figure 5 and Table 1.

**Figure 4.**
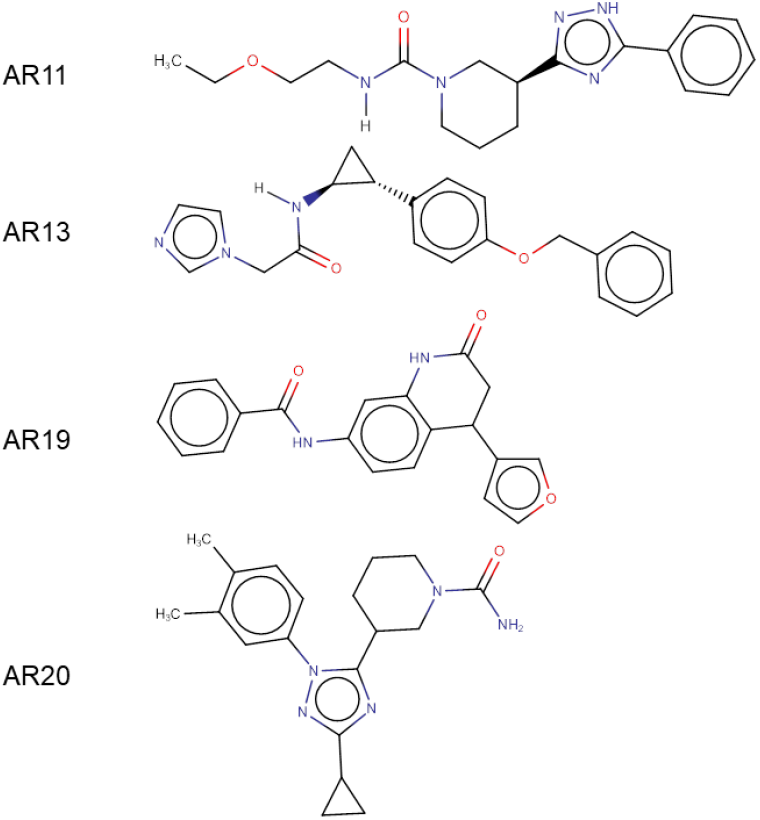
Four most active aromatase inhibitors from virtual screening.

**Figure 5.**
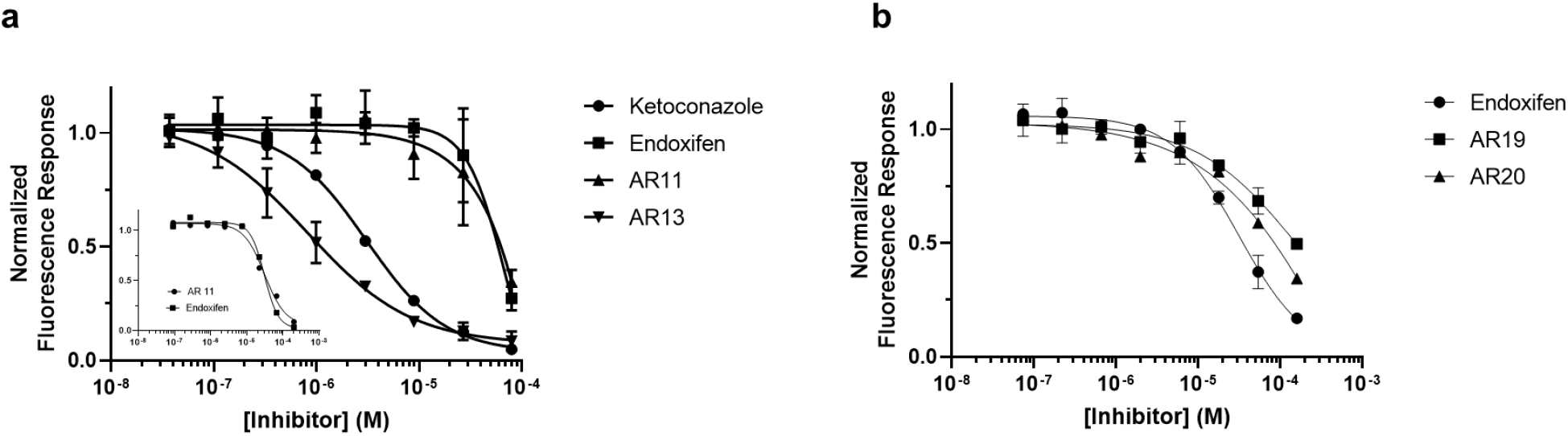
Dose-response curves at 7.5 nM P450 and 25 μM MFC substrate with varying concentrations of inhibitors a) AR11 and AR13, and b) AR19 and AR20. Semi-log curves are fit to a 4-parameter logistic model with data from positive control inhibitors ketoconazole and endoxifen. Reaction mixtures with a 3-fold increase in concentration for AR11 and endoxifen are provided as an inset in (a).

**Table 1.** Activity data collected from active compounds fit to a 4-parameter logistic model.

| *Inhibitor* | *IC<sub>50</sub> (μM)* | *Hill slope* | *Fractional activity at saturation* | *Goodness of fit ($R^2$)* |
| --- | --- | --- | --- | --- |
| *ketoconazole* | 3.08 | 1.13 | 0.035 | 0.999 |
| *endoxifen* | 30.69 | 2.26 | 0.021 | 0.997 |
| *AR11* | 31.07 | 1.51 | 0.051 | 0.986 |
| *AR13* | 0.82 | 0.86 | 0.071 | 0.978 |
| *AR19* * | 65.87 | n/a | 0.144 | 0.845 |
| *AR20* | 42.83 | 0.86 | 0.028 | 0.992 |

\*3-parameter model reported with a Hill coefficient of 1.

**Table 2.** Steady-state kinetic constants in the presence of inhibitors AR11 and AR13 at 10 nM P450.

| | $V_{\max}$ (pmol<br>HFC/min/pmol P450) | $K_m$ (μM) | $K_i$ (μM) | $\alpha$ | $K_i'$ (μM) |
| --- | --- | --- | --- | --- | --- |
| No Inhibitor | 0.689 | 24.38 | - | - | - |
| AR11 | 0.678 | 24.60 | 14.920 | 2.567 | 38.300 |
| AR13 | 0.654 | 22.01 | 0.039 | - | - |

### Optical absorption properties of active compounds

P450s exhibit signature Soret peaks that are detectable in the 390 – 460 nm range ^39^. Perturbations to the heme environment yield absorption shifts that are conserved. In the absence of substrate, water typically occupies the sixth site of the iron octahedral complex. This is evidenced by a Soret peak typically in the 415 – 417 nm range ^39^. Interactions between the heme iron and stronger field ligands will induce absorption shifts at longer wavelengths. The addition of the native substrate, androstenedione (ASD), causes water displacement due to the C-19 protrusion from the ASD backbone ^40^. This is indicated by a hypsochromic shift typically in the 390 – 394 nm range due to iron’s adoption of a 5-coordinate high-spin state (5-CHS) ^39^. We report that the heme cofactor in recombinant Cyp19 displays a Soret peak shift in 100 mM PPB (pH 7.4) at 416 nm (blue trace) and 395 nm in the absence and presence of ASD, respectively. Titration of AR13 induces an 8 nm bathochromic shift to 424 nm in the absence of ASD (Figure 6a). This indicates an interaction with the iron by a stronger field ligand than water. Additionally, the enzyme may reversibly adopt the 5-CHS in the presence of AR13 with a 25:1 molar excess of ASD to inhibitor (Figure 6b). Further, the Cyp19-AR13 adduct can be reduced with dithionite to bind CO reversibly. The subsequent addition of AR13 fully recovers the 424 nm peak after 1 hour.

**Figure 6.**
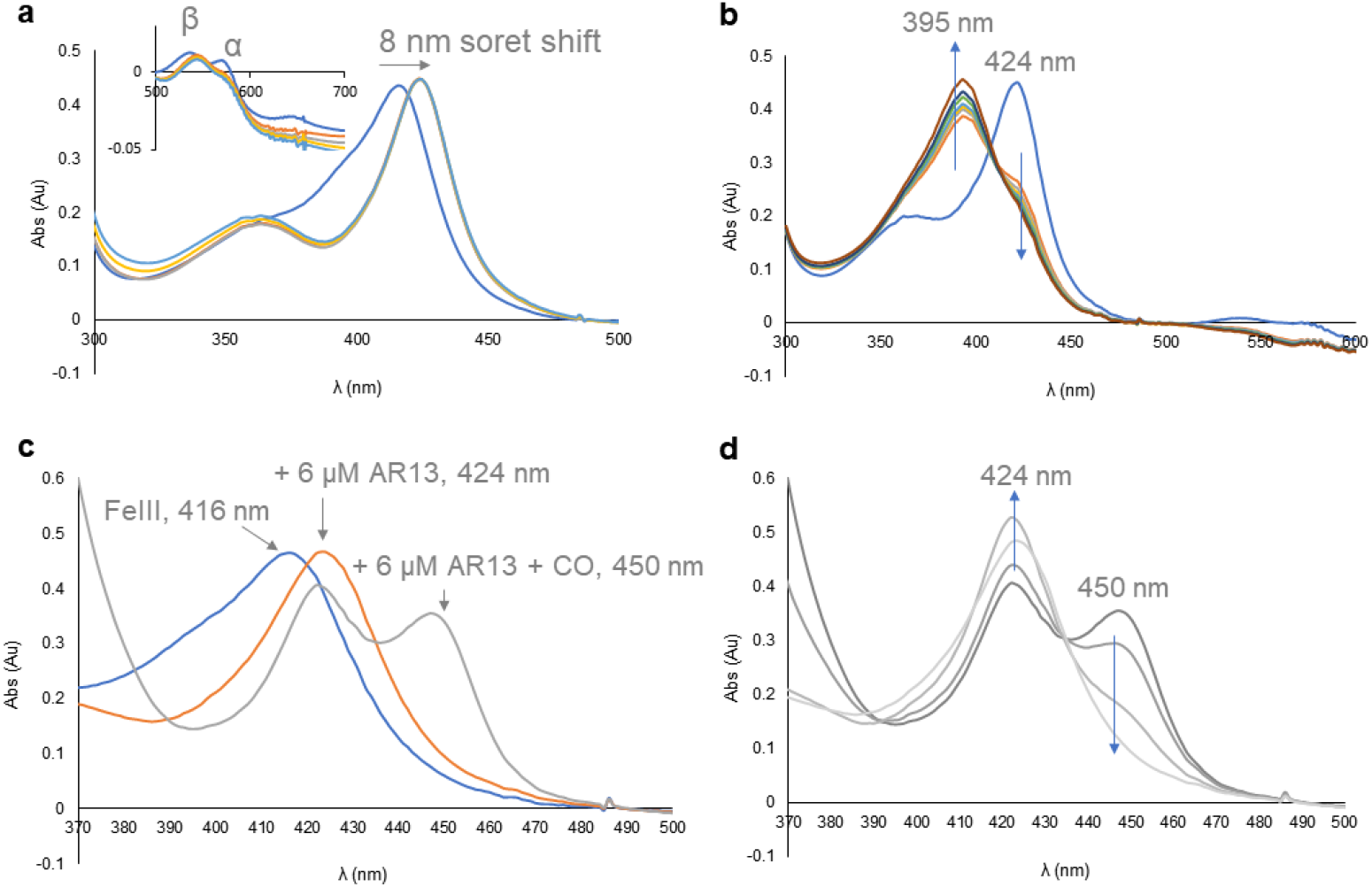
Soret peak shifts in the presence of inhibitor AR13. a) Bathochromic shift of Soret peak maximum from 416 to 424 nm upon titration with AR13 at 6, 12, 60, and 120 μM AR13. Inset shows a 5 nm shift in the β band from 535 to 545 nm and loss of the α band at 570 nm. b) Time-dependent increase of P450 in the high-spin state after the addition of 70 μM ASD to a reaction mixture with 2.5 μM AR13. c) P450 peak is observed after dithionite and CO addition to a reaction mixture of 3 μM P450 and 6 μM AR13. Subsequent addition of 6 μM AR13 results in the recovery of the 424 nm Soret peak and complete loss of CO-bound enzyme in a time-dependent manner (d).

Titration of up to 100 μM of endoxifen did not induce a Soret peak shift from 395 nm in the Cyp19 ASD-bound state (Figure 7a). This indicates that ASD remains in the active site in the presence of a 50-fold molar excess of endoxifen. This behavior is expected of a noncompetitive inhibitor where the K_i_ is unchanged in the presence of substrate. At a 3-fold (6 μM) molar excess, AR11 prompted the appearance of a peak shoulder near 416 nm. Figure 7b illustrates the gradual increase of the peak shoulder at 416 nm with an increase in the concentration of inhibitor. This indicates that AR11 causes the enzyme to favor a shift to the 6-coordinate low-spin state and the displacement of ASD in the active site. At 100 μM AR11, the inhibitor-bound 6-coordinate low-spin state is apparent at 416 nm (Figure 7d). The inset in Figure 7d illustrates that AR11 causes Cyp19 to favor the low-spin state in the absence of its native substrate.

**Figure 7.**
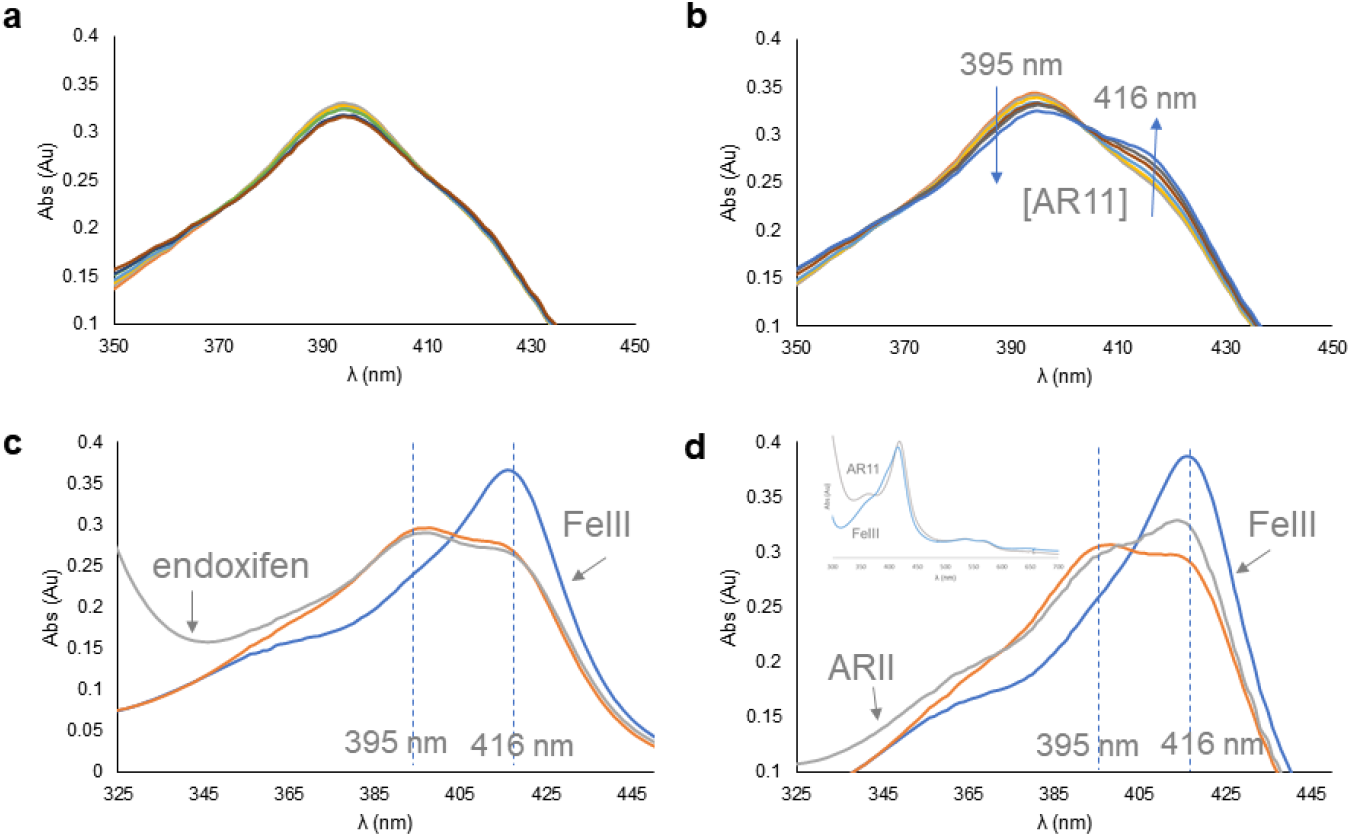
Optical absorption spectra of reaction mixtures in the presence of 3 μM P450, 2 μM ASD, and various concentrations of endoxifen (a and c), and AR11 (b and d). Titration with inhibitor from 6 – 36 μM (a – b), and at 100 μM (c – d). The addition of AR11 favors a transition to the 6-coordinate low spin state indicated by a hyperchromic shift at 416 nm. Blue arrows in panel b indicate the absorption trend as the concentration of AR11 increases. Inset in panel d shows Cyp19 in the absence and presence of 100 μM AR11.

AR19 and AR20 share the same Soret peak trends in the presence and absence of 2 μM ASD. Both compounds induce a gradual shift to the iron low-spin state indicated by simultaneous hyperchromic and red shifting towards 420 nm. This indicates that the population of Cyp19 with iron in the 6-coordinate state increases in the presence of inhibitor. The red shift from 416 nm (blue trace) to 420 nm (red trace) is apparent in Figures 8c and 8d. At 100 μM AR19, 2 μM ASD introduces a faint peak shoulder at 395 nm. Although there is a drop in the absorption at 420 nm, a peak shoulder is not apparent at this concentration. Higher concentrations of AR19 are required to induce the same Soret peak effects as AR20, suggesting that AR20 is a more potent inhibitor.

**Figure 8.**
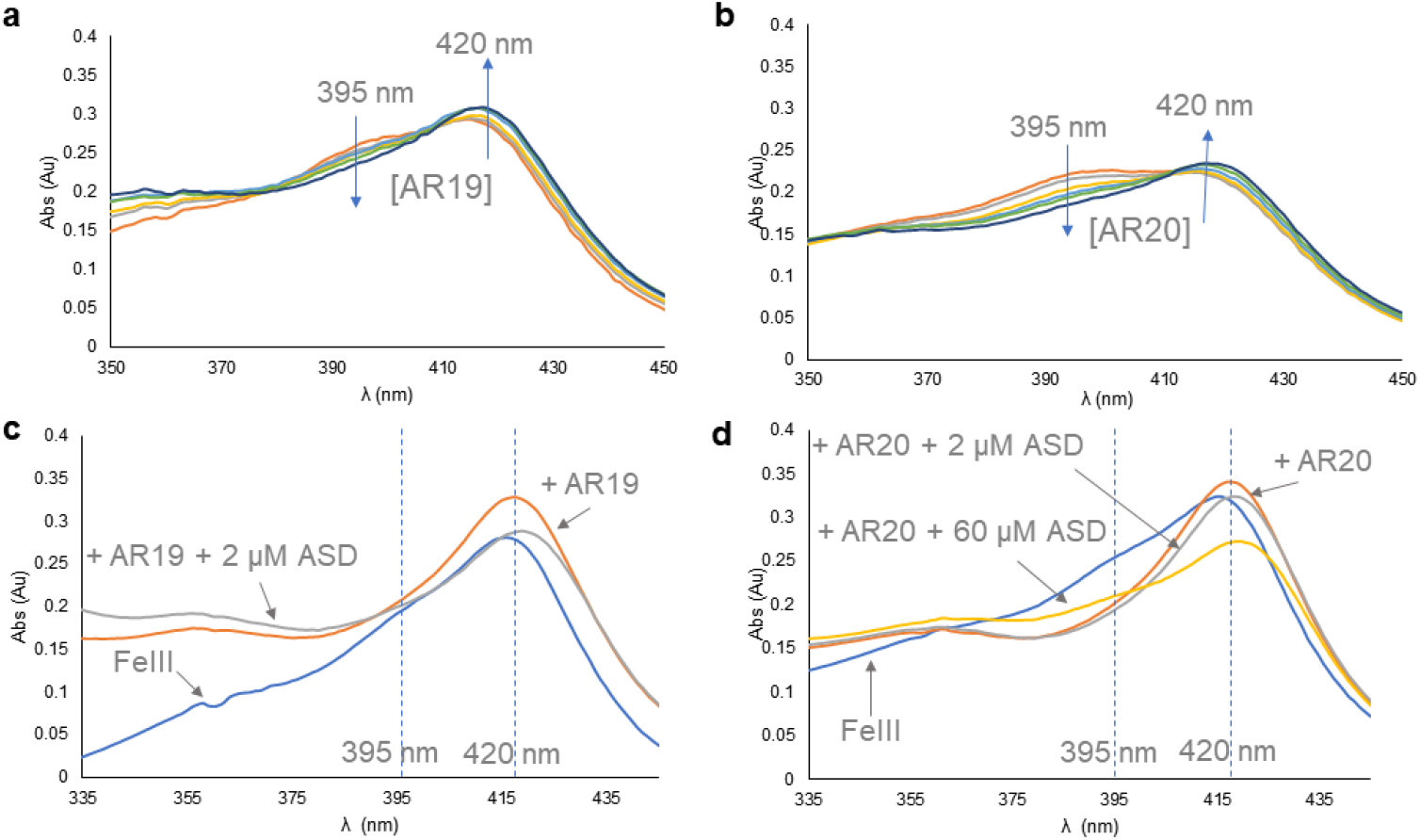
Optical absorption spectra of reaction mixtures in the presence of 2 μM P450, 2 μM androstenedione (ASD), and various concentrations of AR19 (a and c), and AR20 (b and d). a-b) Titration with inhibitor 6 – 36 μM favors a transition to the 6-coordinate low spin state. Blue arrows indicate the absorption trends as the concentration of inhibitor increases. c – d) Addition of 100 μM inhibitor to Cyp19 induces a 4 nm shift from 416 nm to 420 nm. A 395 nm peak shoulder is apparent after the subsequent addition of the native substrate, ASD.

### AR11 and AR13 steady-state kinetic assays

Kinetic assays of our two most potent inhibitors were performed under steady-state conditions to determine their Michaelis-Menten and inhibition constants (Figures S4, S5). These assays were used as a diagnostic tool to confirm whether AR11 and AR13 interact at the active site or a distinct site. Inhibition assays were performed at various concentrations of MFC up to 2 X K_m_. Nonlinear regression analyses in the absence of inhibitor yielded substrate MFC V_max_ and K_m_ values of 0.689 pmol HFC/min/pmol P450 and 24.38 μM.

Lineweaver-Burk plots of AR11 yielded functions that intersected the y-axis at different inhibitor concentrations. These corresponded to different apparent V_max_ values, suggesting that both inhibitors do not act competitively. Visual inspection of Dixon-type plots also suggested that AR11 reduces the V_max_. In Figure 9, the intersection of regression curves in panel A corresponds to the K_i_ value. The intersection of the curves in panel C corresponds to a K_i_ ′ or inhibitor dissociation constant in the presence of substrate MFC. This value infers non-mutual exclusive binding. As such, AR11 was fit to a mixed-type nonlinear function.

**Figure 9.**
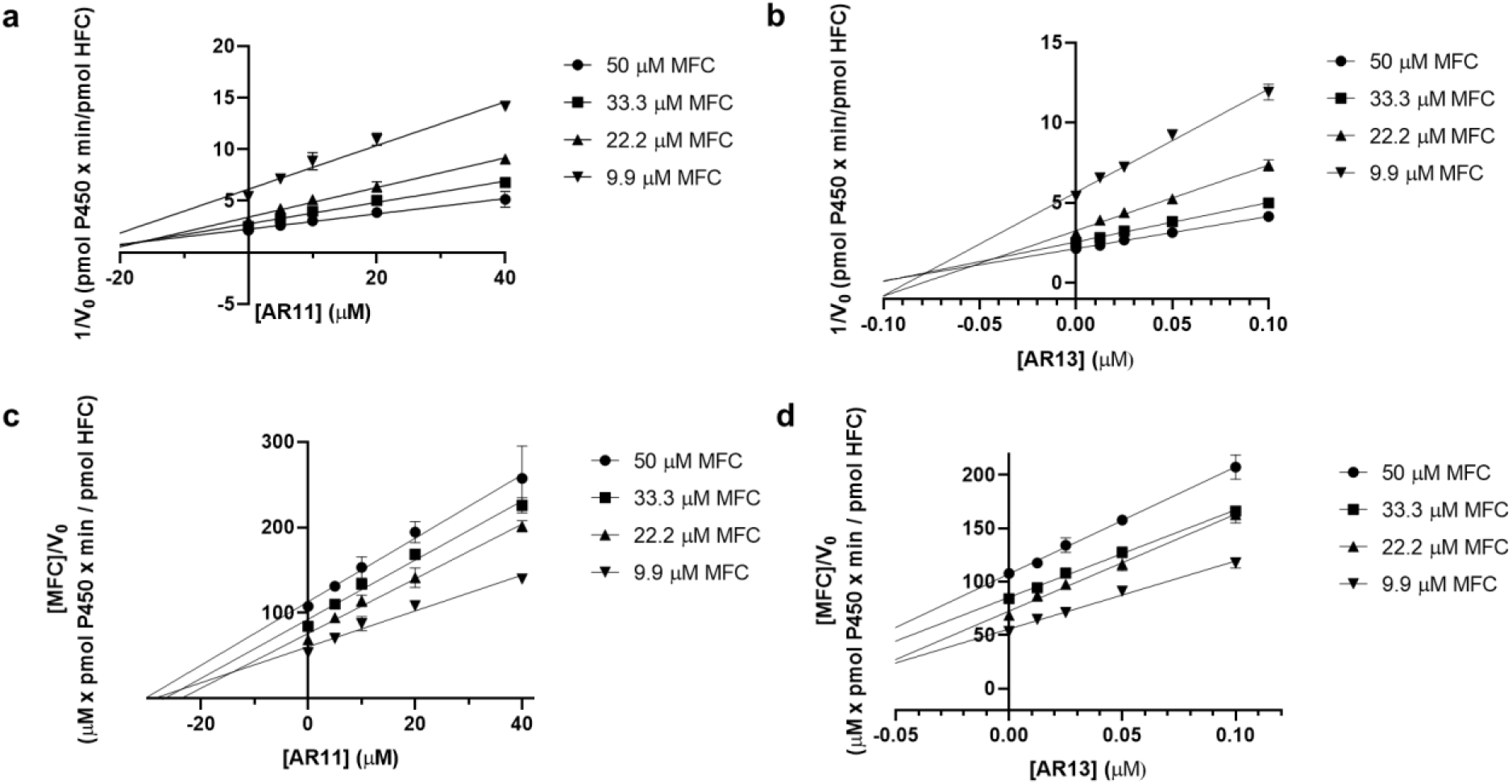
Dixon-type inhibition plots for AR11 (a,c) and AR13 (b,d).

Lineweaver-Burk analyses of MFC conversion at different concentrations of AR13 inferred nonmutual exclusivity between substrate MFC and AR13 since the apparent V_max_ varied with inhibitor concentration. On the contrary, Dixon-type plots suggested that MFC and AR13 are mutually exclusive since the K_i_’ value was indeterminate (Figure 9d). Nonlinear regression fit to a competitive-type function yields a K_i_ of 39 nM, 35 – 44 nM for a 95 % confidence interval. This is within an order of magnitude from the working enzyme concentration and supports tight-binding behavior. In a Lineweaver-Burk plot, the 1/V_0_ values of tight-binders begin to converge at high substrate concentrations. Curvature in the AR13 data points is most recognizable at high inhibitor concentrations.

Non-linear regression for AR11 and AR13 fit to their diagnosed mode of inhibition returned K_m_ values of 24.60 and 22.01 μM at 10 nM P450. Table S4 presents the effects of each inhibitor on these kinetic constants. The apparent K_m_ increases in the presence of each inhibitor, indicating that more substrate MFC is needed to achieve ½ V_max_. Meanwhile, the maximum velocity only decreased with AR11. This indicates that MFC saturation will not attain full enzyme activity in the presence of AR11. Therefore, both substrate and AR11 bind Cyp19 at discrete sites. In contrast, the ability to achieve the V_max_ with a molar excess of MFC over AR13 indicates that both compounds bind the active site.

### MD simulations of compounds docked against Cyp19

Molecular dynamics simulations were performed to observe the residence time of the inhibitors in the proximal heme binding pocket. All our active compounds, except AR11, dissociated from the proximal heme site in less than 1 μs suggesting modest binding strength. AR13 exhibited the weakest interaction at the proximal heme site. It remained in the pocket for less than 100 ns in all replicates.

Since the Soret shifts supported an iron-imidazole interaction, AR13 was docked to the catalytic site in 25 independent runs. The binding mode of each enantiomer yielding the closest iron- imidazole distance was used for MD simulations. They bound with free energies of −36 ± 4 kcal/mol (1R,2S) and −36 ± 2 kcal/mol (1S,2R) as calculated by MM-PBSA. Both enantiomers are projected to form a π-π interaction with F221 in the substrate access channel and a low energy π-π interaction between the terminal azole and the heme’s porphyrin system. The Fe-N distances for each compound averaged 6 Å, indicating an indirect heme interaction.

More rigorous global redocking of AR11 located it in the substrate access channel more often than in the proximal heme site. AR11 formed π-π interactions with F221 and W224 and a hydrogen bond between the triazole and D309. This suggests that the triazole partially obstructs a space occupied by ASD since E309 is involved in a critical contact with the C3-carbonyl of androgens. Production runs yielded an average free energy of −29 ± 8 kcal/mol. In the proximal heme site, the free energy was −32 ± 4 kcal/mol, despite a destabilizing effect from E357. We report the putative binding modes at both the proximal heme site and substrate access channel in Figure 10 since their binding energies have overlapping confidence intervals. We include the decomposition scores of the interactions with the greatest contributions to these energies.

**Figure 10.**
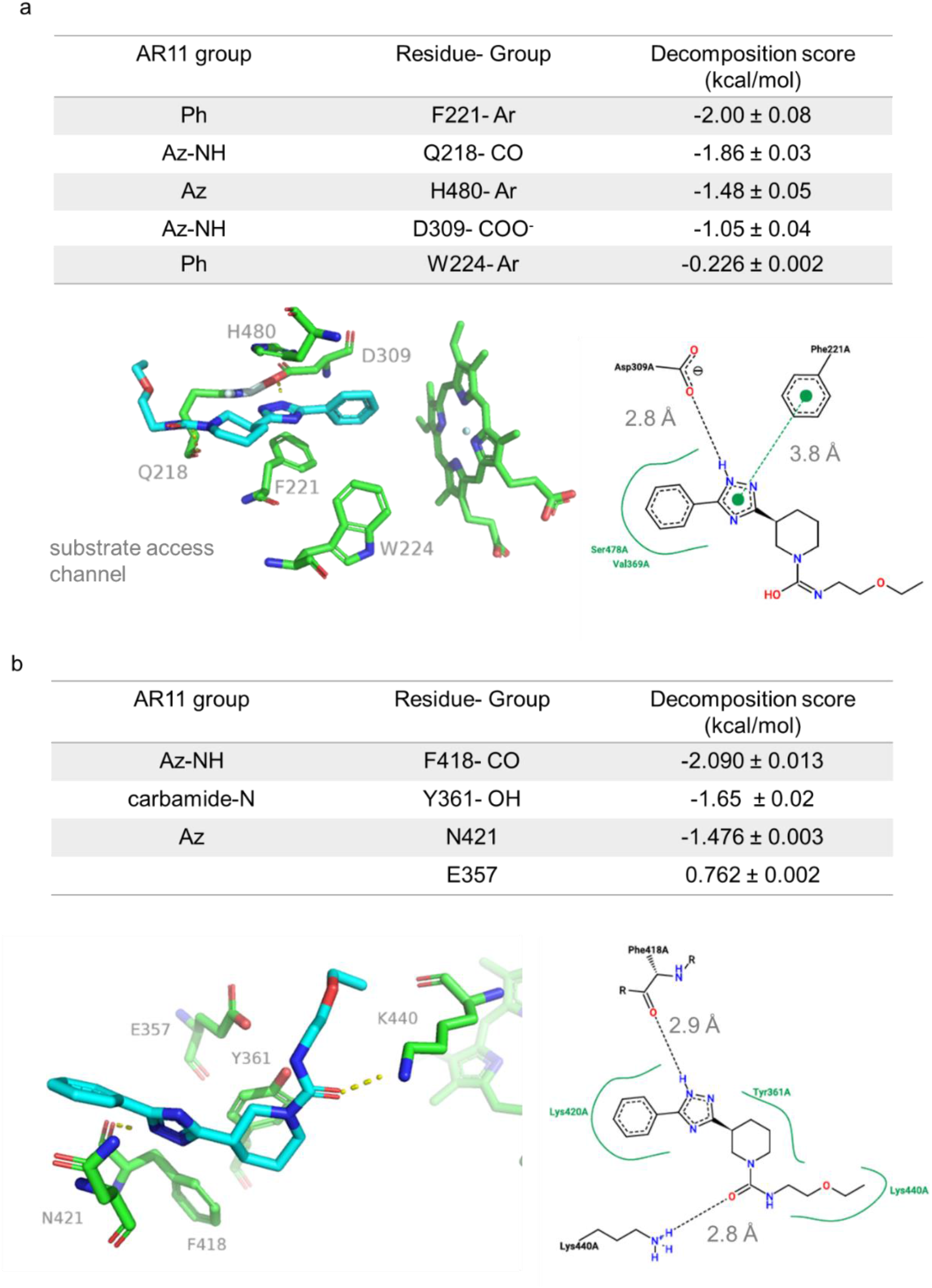
3-D and 2-D interaction diagrams for AR11 with decomposition scores of the projected protein-ligand interactions at the substrate access channel (a) and the proximal heme site (b). 2-D depictions were generated with PoseView software ^41^. Az- azole, Ph- phenyl, Ar- aromatic groups.

**Table 3.** Docking and simulation metrics of endoxifen and novel Cyp19 inhibitors.

| *Ligand* | *Proximal heme site average dissociation time (ns)* | *Proximal heme site average binding free energy (kcal/mol)* | *Substrate access average binding free energy (kcal/mol)* | *Active site average binding free energy (kcal/mol)* |
| --- | --- | --- | --- | --- |
| *E-end* | < 200 | -17 ± 4 | - | - |
| *Z- end* | < 500 | -24 ± 3 | - | - |
| *AR11* | >1000 | -32 ± 4 | -29 ± 8 | - |
| *AR13 (1R,2S)* | <100 | NR | - | -36 ± 4 |
| *AR13 (1S,2R)* | - | - | - | -36 ± 2 |
| *AR19* | 400** | NR | -33 ± 3 | - |
| *AR20* | 800* | -21.92 ± 0.04* | - | - |

\*denotes a value from 1 replicate run
\*\* denotes a value from 2 replicate runs

AR19 and AR20 dock to the proximal heme site but dissociate after developing unfavorable interactions with E357, as demonstrated by positive decomposition scores (Figures S6, S7). We also note that control inhibitors, E- and Z-endoxifen, disassociate due to positive free energy contributions from E357. In 1 out of 3 simulations, AR20 re-associates and remains in the site > 1 μs with −22 kcal/mol average binding free energy (Figure S8). We do not report decomposition scores for AR20 because it only transiently remains in a single binding mode throughout the production run. AR19 formed stable interactions in the substrate access channel with F221 and E309. Further, it had a proclivity to migrate closer into the active site for the duration of the production runs (Figure 11a). The average free energy was −33 kcal/mol.

**Figure 11.**
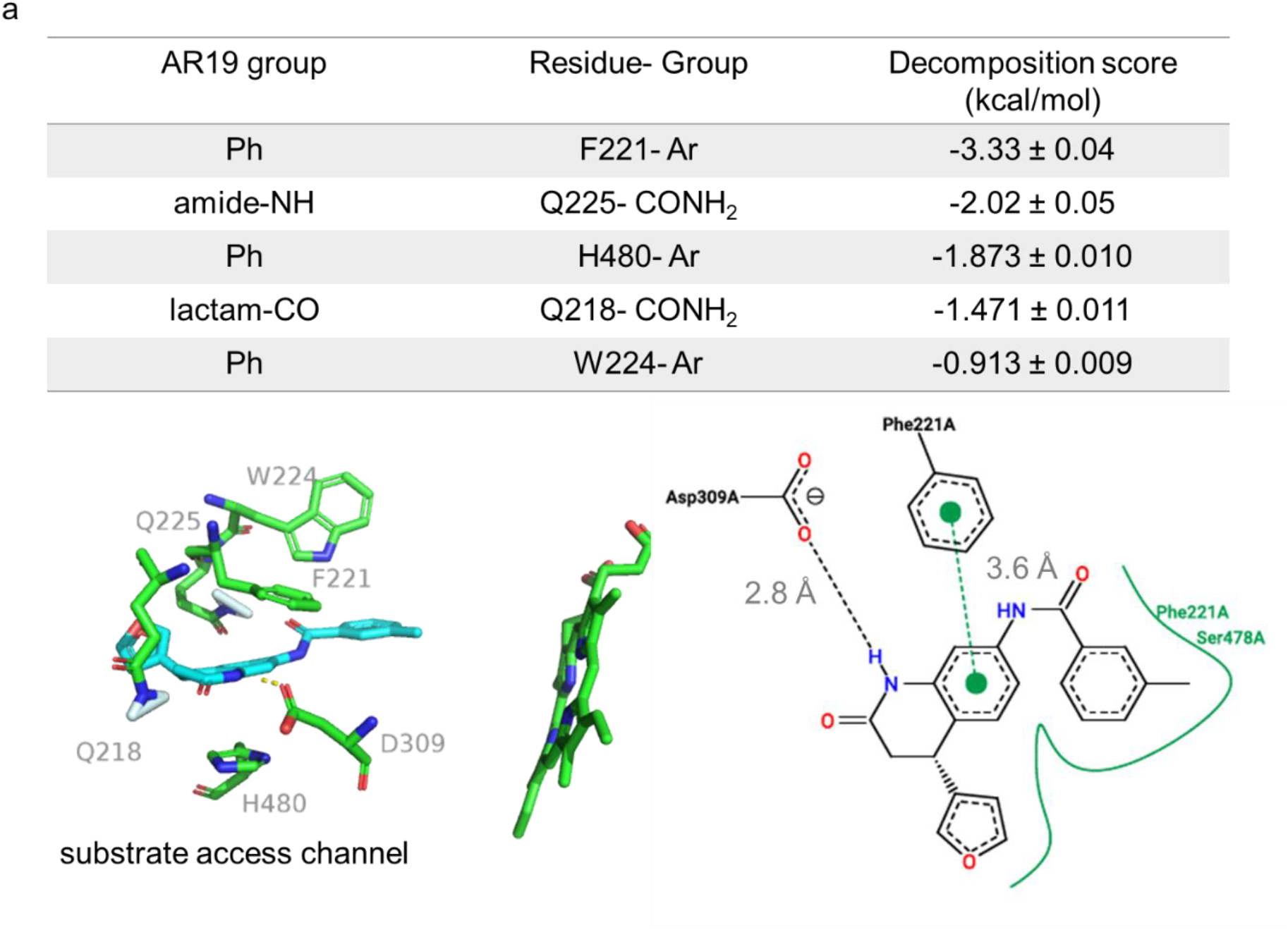
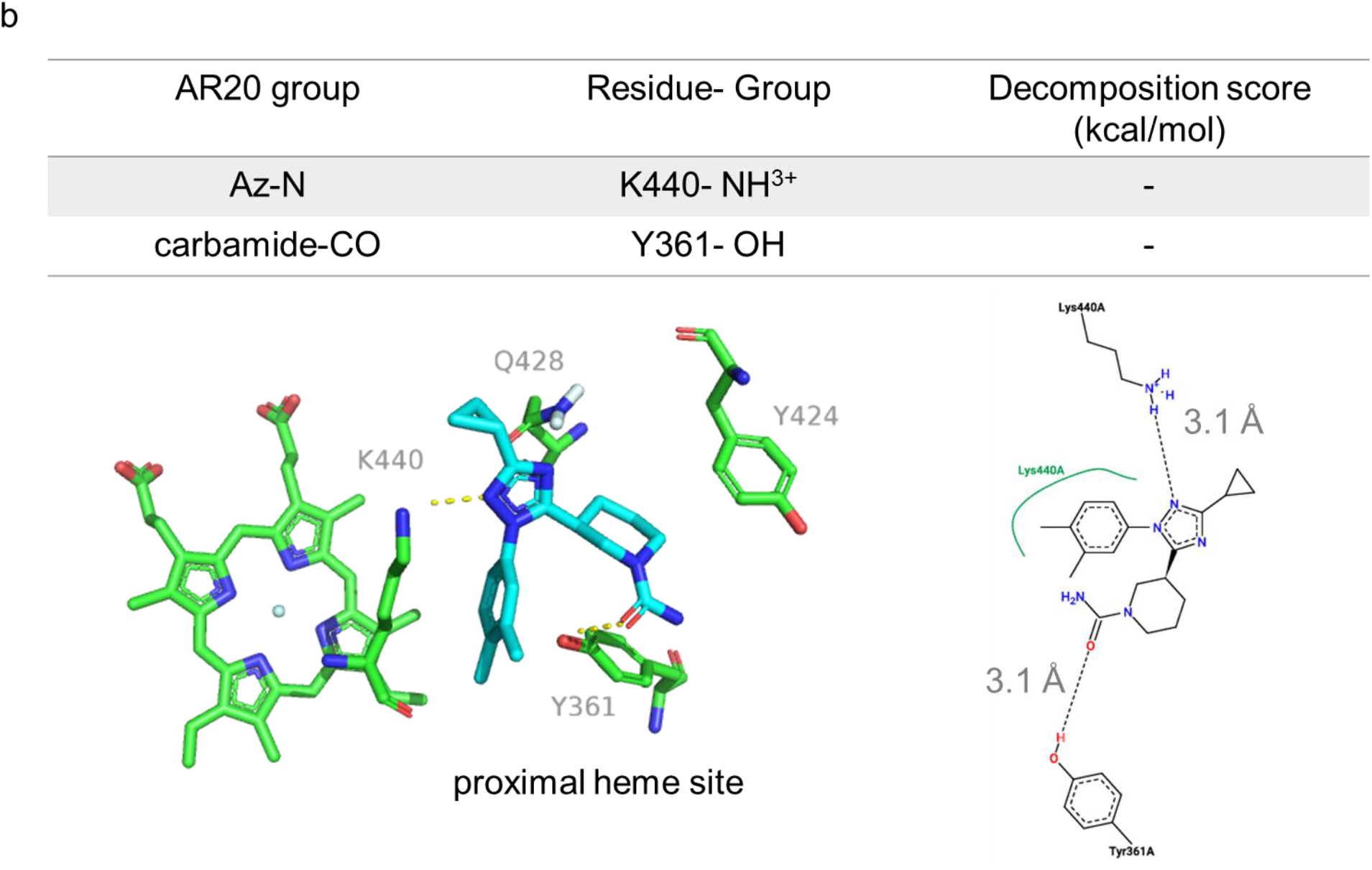
3-D and 2-D interaction diagrams with decomposition scores of the projected protein-ligand interactions for AR19 (a) and AR20 (b). 2-D depictions were generated with PoseView software ^41^. AR20 binding mode represented here is for replicate 2, and >280 ns. Decomposition scores are not available for AR20. Az- azole, Ph- phenyl, Ar- aromatic sidechain.

## Discussion

MD simulations support that none of the inhibitors identified bind stably at the Cyp450 reductase binding site. Rather, AR13 binds the enzymatic active site, as our inhibition kinetics data demonstrates, correlating well with the absorption data we present. The 8 nm Soret peak shift in the presence of AR13 indicates an interaction with the heme iron in the active site. Strong red shifts from a water-bound heme typify occupation of a strong, nitrogenous sigma-donor ligand. This is likely an interaction with AR13’s terminal imidazole moiety. A direct Fe-N interaction was confounded by our MD simulations that predicts a 6 Å distance. This would indicate a water-bridged state. Therein lies the possibility that there could be multiple energetically favorable 6-coordinate states in equilibrium. Although there are reports of P450-azole ternary complexes, the compounds under study appear to be limited to triazoles that induce weaker soret peak shifts. Incontrovertible evidence warrants further study.

Cyp19’s ability to bind ASD and CO in the presence of AR13 demonstrates its capacity to maintain its functionality. Above all, the need for a 28-fold molar excess of ASD to induce a blue shift to 395 nm supports the inhibitor’s high potency. Among all our inhibitors, AR13 was the most potent. This correlates with its predicted average binding energy being the greatest over AR11, AR19, AR20, and endoxifen. AR13 has a 40-fold lower IC_50_ value than that of endoxifen and interacts with Cyp19 on the same order of magnitude as norendoxifen (K_i_ = 35 nM). Furthermore, its chemical structure is remarkably distinct from current third generation NSAIs. To this end, AR13 may be useful as a scaffold to design new NSAIs.

Titration with AR11 against androstenedione induces a peak shoulder between 414 – 416 nm, indicating that AR11 displaces ASD while substrate access to the heme iron is maintained. The average binding free energies at the proximal heme site and substrate access channel have overlapping confidence intervals and the Hill coefficient from a dose-response curve is greater than 1.5. As such, we project that both sites presented in this work are occupied at saturating concentrations of AR11. Likely, AR11 preferentially interacts at the substrate access channel since E357 is projected to be destabilizing. The transition to a 5-coordinate out of-plane position may cause the observed 2.6-fold increase in the K_i_’ at the proximal heme site. However, there are currently no crystal structure of Cyp19 in the 6-coordinate water-bound state for comparison.

We note that our control inhibitor, E/Z- endoxifen, exhibits similar trends as AR11 in this study. Both dock to the proximal heme site and consistently yield a Hill coefficient close to 2 suggesting multiple binding sites. We note two important differences. Firstly, MD simulations predict that E- and Z- endoxifen do not remain in the proximal heme site, which is consistent with work by Sgrignani et al ^42^. Secondly, E/Z-endoxifen does not cause dissociation of ASD from the active site unlike AR11. This is potentially due to AR11 entering a pocket that is partially occupied by ASD.

Titration with AR19 and AR20 causes a 4 nm red shift from a water-bound iron to 420 nm. AR20 is a 1,2,4-triazole that may induce such a shift if there is a direct interaction with the heme iron or formation of a water-bridged ternary complex. P450 ternary complexes have been reported with 1,2,3- and 1,2,4- triazoles ^43^. Although this suggests that AR19 and AR20 bind similarly to the active site, small Soret peak shifts may also arise from a change in the iron’s chemical environment. Heme perturbation induced by allosteric interactions may cause such Soret peak shifts.

Altogether, we conclude that E/Z- endoxifen, AR11, and AR19 preferentially occupy the substrate access channel, inhibiting Cyp19 on the same order of magnitude. Since the substrate access channel is dynamic, compounds forming weaker interactions such as AR19 may migrate towards the active site to form a direct interaction with iron. E/Z- endoxifen may bind the substrate access channel with high affinity and a second site at saturating levels of inhibitor. We expect that AR11 interacts at the substrate access channel and the proximal heme site at saturating levels. In the substrate access channel, AR11 can bind deeper into the pocket such that it partially occupies space by ASD near the catalytically active E309. AR20 is projected to transiently interact at the proximal heme site.

The proximal heme site is acting as a low-affinity binding site for multiple inhibitors. The proximal heme site is an attractive target to modulate P450 activity because it is a conserved feature among P450s and distinct from the catalytic core. Despite class 2 P450s having less than a 40% sequence identity, all of them are expected to partner with CPR at this site. CYP19 may serve as a model to study its druggability for three reasons. Firstly, a crystal structure with the pentameric PEG bound at the proximal heme site is available and PEG inhibited enzyme activity in a time-dependent manner ^44^. Secondly, Cyp19 has a well-defined cavity of 584 Å^3^, roughly 200 Å^3^ larger than the active site, that can accommodate a larger library of compounds ^44^. Lastly, the loop region between helices k” and L contains the meander region of 21 residues long. Roughly 30 – 40% of P450s have a loop 14 or 15 residues long- all of which are class 2 P450 enzymes. This disparity offers a niche to selectively target CYP19 as less than 5% of P450s have a meander loop as lengthy ^45^. It is likely that higher affinity inhibitors to the proximal heme site can be identified.

Other P450s that may be allosterically inhibited at the proximal heme site include Cyp3A4 and Cyp1A2. They are the major enzymes in drug metabolism and have well-documented heterotrophic effectors ^46–48^. It is possible that these P450s can be modulated at the proximal heme site as the cavities are large enough to accommodate ligand-binding. The Cyp3A4 and Cyp1A2 pockets are 1.1 and 0.7 times the size of the Cyp19 proximal cavity ^44^. More research is needed to investigate alternative ligand binding sites in P450s ^49–52^.

In a broader context, nearly all exogenous compounds are subject to redox chemistry by P450 enzymes. Azoles and furan-containing xenobiotics, such as those presented here, are often P450 substrates ^53^. There are also many important azole-containing drugs that are used to treat fungal infections ^54^. However, they also inhibit human P450 isozymes and undergo various metabolic reactions. Off-target interaction with P450s may lead to toxicity or adverse drug interactions.

## Acknowledgments

We thank Wesley Yoshida (Univ. Hawaii), Joshua Gurr (Univ. Hawaii), and the Kansas State University Biomolecular NMR Lab for the analytical work on compounds AR11 and AR13. In addition, we acknowledge the MERCURY Consortium for use of Skylight in the MD simulations presented in this work. We thank F. Peter Guengerich (Vanderbilt University) for providing the aromatase expression plasmid and general technical advice. HN was funded by the NSF CAREER Award 1833181, the Victoria and Bradley Geist Foundation, and the Hawaii Community Foundation.

## Conflicts of Interest

The authors declare no competing interests.

## Notes

### Competing Interest Statement

The authors have declared no competing interest.

