## Supplementary figures and data for "Mechanisms of allosteric and mixed mode aromatase inhibitors"

This file contains Tables S1 – S4, Figures S1 – S9, and detailed methods to produce and characterize functional recombinant aromatase.

Table S1. Residues selected as part of the interaction restraints for prediction-driven docking.

| Cytochrome P450 | Active residues selected for prediction-driven docking | Passive residues selected for prediction-driven docking |
| --- | --- | --- |
| CPR (3FJ0) | 117, 119, 120, 153, 157, 158 | 118, 122, 123, 131, 154, 155, 156, 159, 160, 162, 165, 166, 184, 186, 187, 188 |
| CYP19A1 | 149, 150, 151, 153, 154, 424, 426, 429, 430, 432, 433, 437, 440, 441 | 101, 104, 105, 108, 109, 145, 146, 147, 155, 156, 157, 158, 159, 202, 276, 281, 361, 364, 422, 423, 425, 434, 438 |

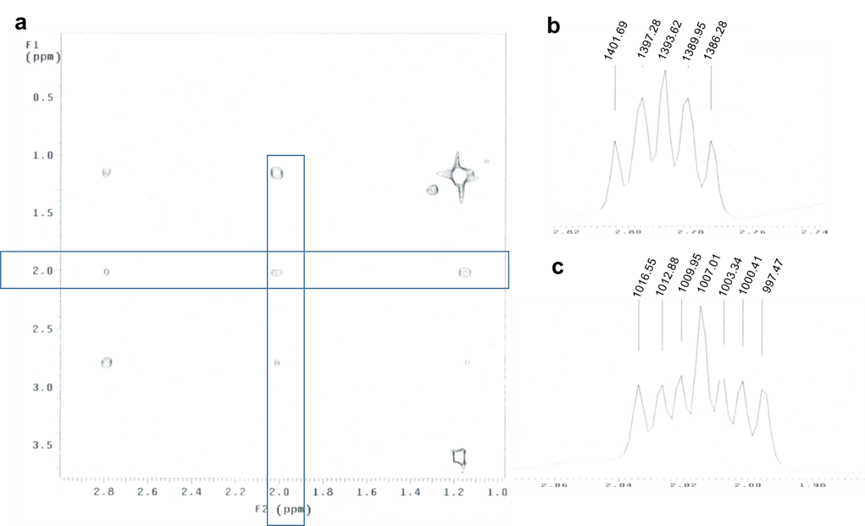

Figure S1. Proton peaks at 1.15, 2.01, and 2.79 at the cyclopropyl group of AR13 in MeOD, and representative ^3^J values indicate geminal hydrogens, and two methine groups with trans hydrogen. a) ^1^H 2-D TOCSY indicate that crosspeaks at 1.15, 2.01, and 2.79 are in the same spin system of the cyclopropyl moiety. b) Peak splitting in hertz at δ 2.79 yield ^3^J values 3.70, 3.70, and 7.97. c) ^3^J values at δ 2.01 are 3.4, 6.5, and 9.6. Coupling constants for the proton at each chemical shift indicate one cis, and two trans neighboring protons. In CDCl_3_ solvent with ^1^H decoupling at δ 2.79, ^3^J values 6.52, and 9.78 Hz remain.

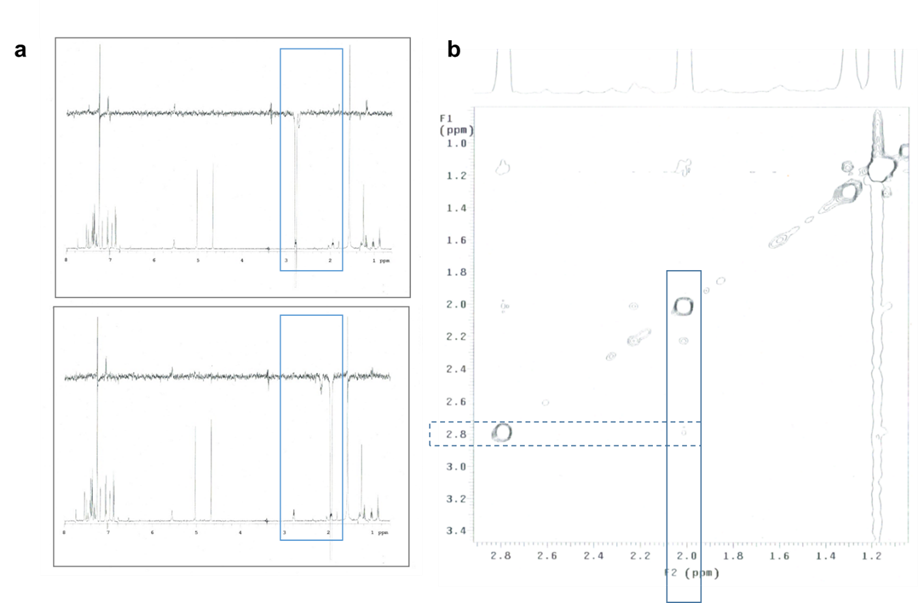

Figure S2. NOEs indicate weak through-space coupling between protons at δ 2.01, and 2.79 on AR13 cyclopropyl group. a) 1D NOEs show no noticeable coupling between protons boxed in blue. Top frame indicates irradiation at δ 2.79, with bottom frame at δ 2.01. b) 2D NOESY has very weak crosspeaks between protons at δ 2.01 and 2.79.

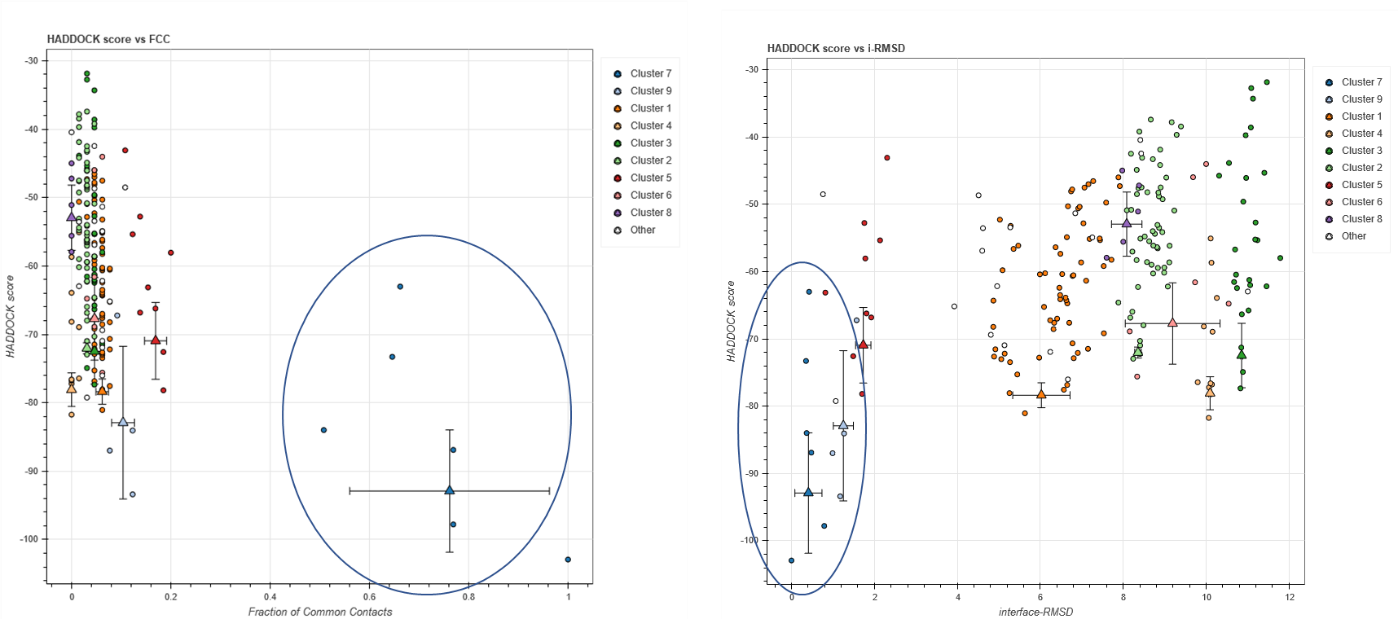

Figure S3. Cluster 7 (circled in blue) returned the most favorable protein-protein interactions as predicted by a lower Haddock score. The left panel indicates that cluster 7 has a greater fraction of common contacts. The right panel indicates lower interfacial RMSD values for members of this cluster.

Table S2. Physicochemical descriptions of the FMN domain–P450 interface for Cyp19 and P450BM3.

| Complex | Interfacial polar/charged interactions | ΔG (kcal/mol)  at 25 ֯C | | K_d_ (M)  at 25 ֯C | K_m_ (M)  experimental | FMN – Heme distance (Å) |
| --- | --- | --- | --- | --- | --- | --- |
| P450BM3 (1BVY) | 17 | -6.9 | 8.6 X 10^-6^ | |  | 22.68 |
| Cyp19-3FJO | 17 | -10.2 | 3.2 X 10^-8^ | | 9.5 X 10^-9 *^ | 14.44 |

*Determined by an estrone-based ELISA by Lo et al. with rat CPR and human P450.^37^

Table S3. Physicochemical properties of chemical compounds.

| Compound | Product no.  (company) | ZINC ID | DSX score | FW (neutral) | logD (pH 7.4) | λ (nm)/ε (M^-1^cm^-1^) |
| --- | --- | --- | --- | --- | --- | --- |
| Androstenedione |  |  |  | 286.41 | 3.93 | 238/14,550 |
| MFC |  |  |  |  |  | 203/-, 334/6,790 |
| HFC |  |  |  |  |  | 203/-, 341/1,891 (391/-, 341 peak shoulder) |
| Ketoconazole |  |  |  |  |  |  |
| Endoxifen |  |  |  |  |  | 243/22,850 282/14,440 |
| AR11 | Z1657610908  (Enamine) |  | -136 | 343.43 | 2.58 | 245/20, 010 |
| AR13 | Z1314885949  (Enamine) | 73608770*  73608774 | -136 | 347.42 | 2.45 | (234/26,090 far UV peak shoulder) 282/4,240 |
| AR19 | 10939549 (ChemBridge) |  | -122 | 346 | 3.04 |  |
| AR20 | 32312044 (ChemBridge) |  | -117 | 339 | 1.75 | 238/9,500 |

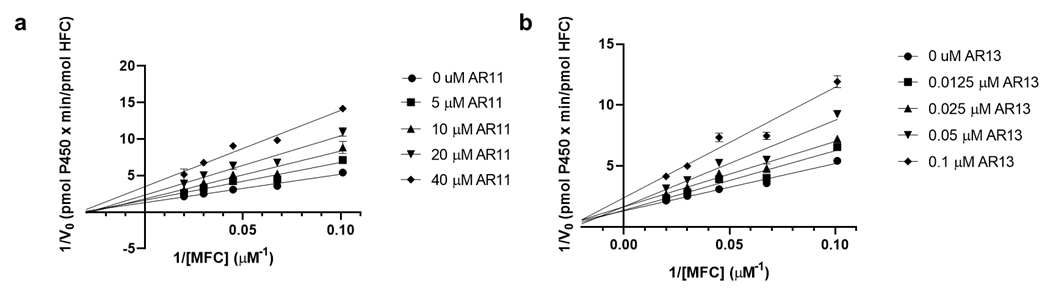

Figure S4. Lineweaver-Burk plots at 10 nM P450 and various concentrations of substrate MFC with inhibitors AR11 (a) and AR13 (b).

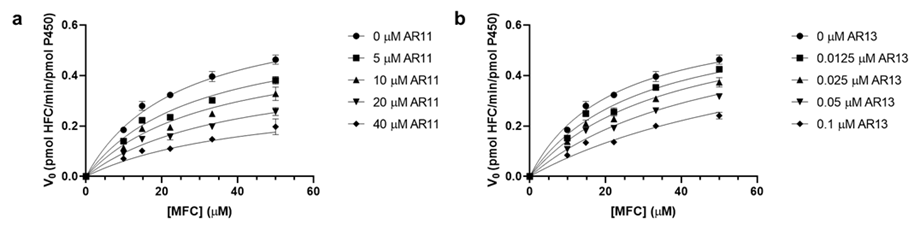

Figure S5. Non-linear regression curves for AR11 (a) and AR13 (b) fit to single-site mixed and competitive-type Michaelis-Menten functions.

Table S4. Apparent Michaelis-Menten kinetic constants at 10 nM P450.

|  | **V_max,app_ (pmol HFC/min/pmol P450)** | **K_m,app_ (µM)** | **R^2^** |
| --- | --- | --- | --- |
| **[AR11] (µM)** |  |  |  |
| 0 | 0.678 | 24.60 | 0.988 |
| 5 | 0.600 | 29.05 | 0.980 |
| 10 | 0.538 | 32.58 | 0.953 |
| 20 | 0.446 | 37.82 | 0.973 |
| 40 | 0.332 | 44.29 | 0.931 |
| **[AR13] (nM)** |  |  |  |
| 0 | 0.654 | 22.01 | 0.989 |
| 0.0125 |  | 29.06 | 0.981 |
| 0.025 |  | 36.12 | 0.984 |
| 0.05 |  | 50.23 | 0.978 |
| 0.1 |  | 78.45 | 0.960 |

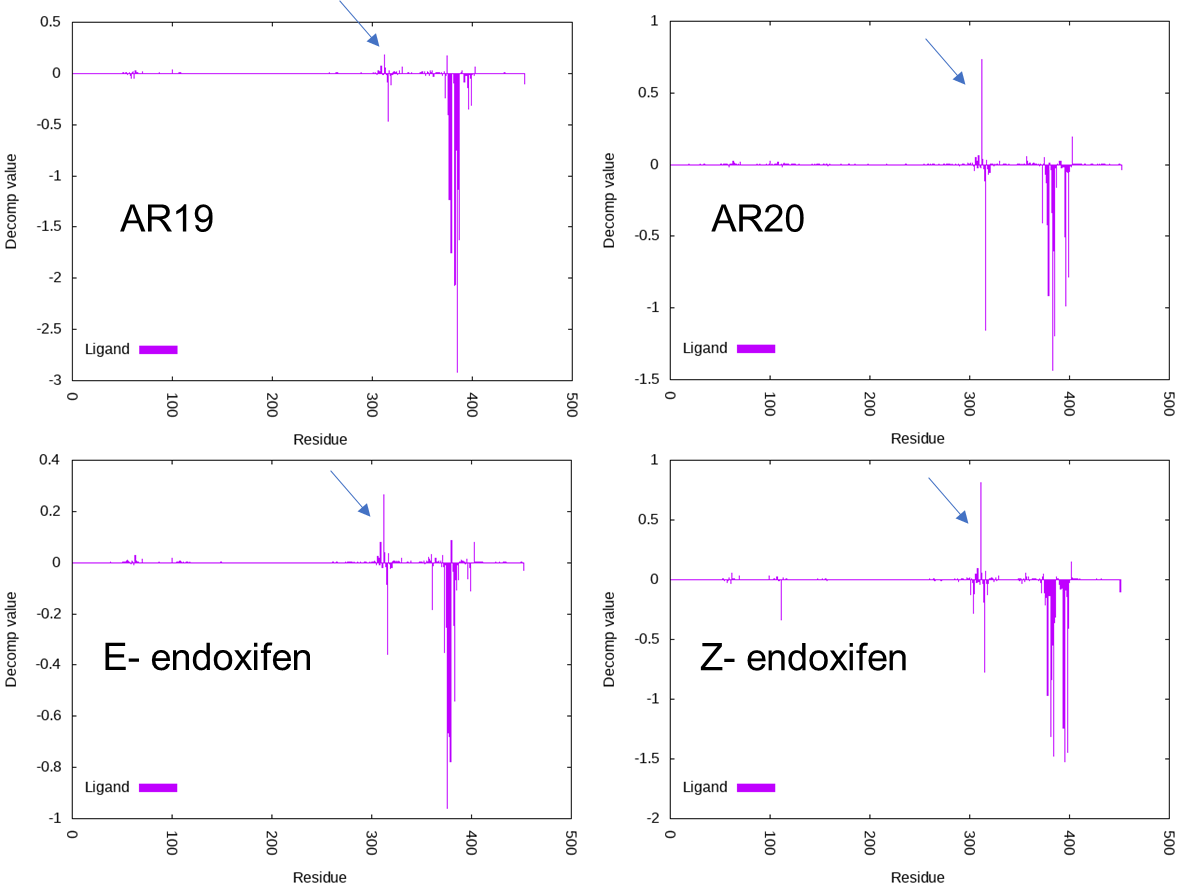

Figure S6. Decomposition values of ligand interaction with each residue (r – 44, r = residue number in wildtype Cyp19) in the proximal heme site. E357 has a positive decomposition value indicative of a destabilizing effect on E/Z-endoxifen, AR19, and AR20 (indicated by a blue arrow).

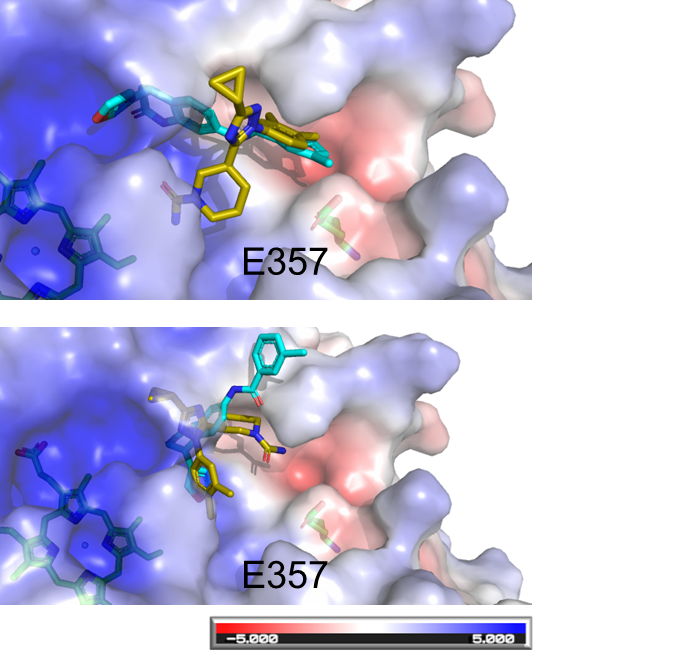

Figure S7. Binding mode of AR19 (cyan) and AR20 (yellow) in the proximal heme site before (top panel) and after (bottom panel) adjustment due to E357 destabilization.

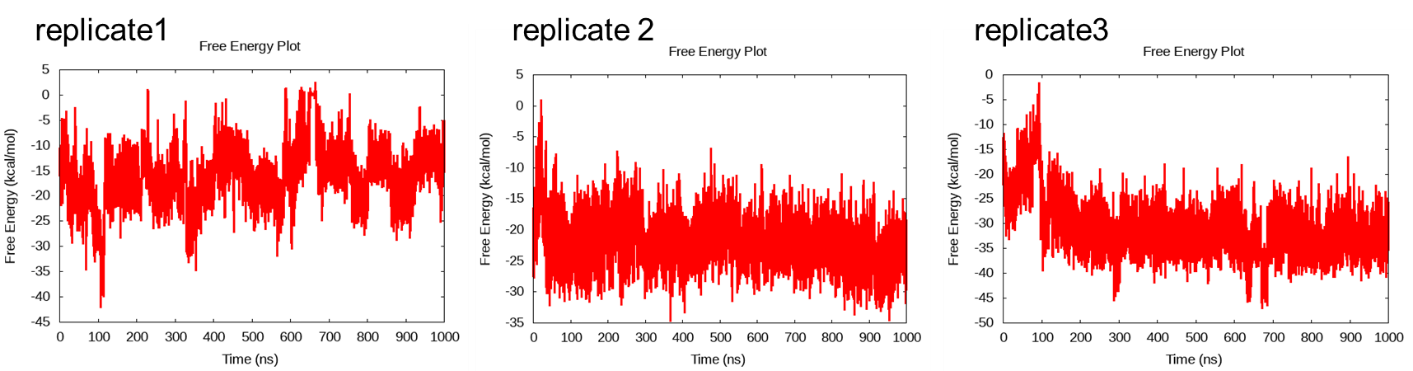

Figure S8. Free energy of binding of AR20 in the proximal heme site. The average free energy we report from replicate 2 remained in the projected Cyp19-CPR interface for the 1 μs run.

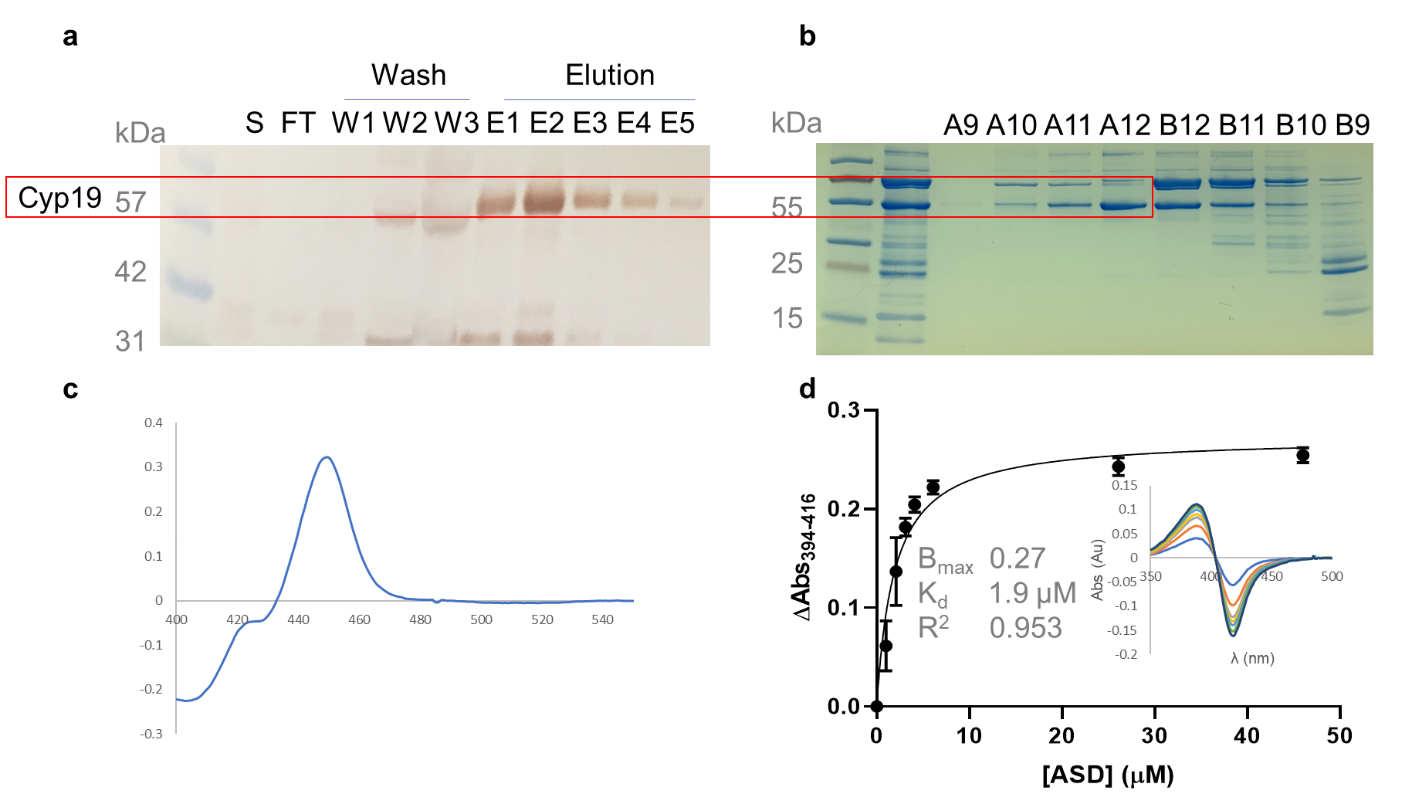

Figure S9. Cyp19 characterization by blot analysis (a), SDS-PAGE (b), CO-difference spectrum (c), and androstenedione binding affinity by absorption difference spectrum (d). a) Blot detection by Ni-HRP of His-tagged Cyp19 with clarified lysate (S), Ni-purified flow through (FT), successive wash steps (W1-3), and successive elution step (E1-5). b) SDS-PAGE of gel filtration fractions with an SEC200 increase column. Lane 2 contains the nickel-affinity purified sample injected onto the SEC column. c) CO-difference spectrum with an internal reference at 490 nm. d) Binding isotherm for Cyp19 single-site binding at 3 μM P450 and titration of ASD with a K_d_ value of 1.9 μM.

**Supplementary Detailed Methods**

**Production, purification, and characterization of Cyp19.** E. cloni 10 G (Lucigen) competent transformants were used for overnight starter cultures in LB Lennox broth and 10 μg/mL carbenicillin at 37 ֯C, 250 rpm. Expression cultures were inoculated with a 1:50 ratio by volume in Terrific Broth media. At OD_600_ = 0.6 – 0.8, cultures were equilibrated to 25 ֯C at 150 rpm for 1 hour prior to induction with 1 mM IPTG and 1 mM δ-aminolevulinic acid. Cultures were harvested and washed with 200 mM NaCl and 100 mM sodium phosphate buffer (pH 7.4).

Cell pellets were resuspended in buffer A (200 mM NaCl, 100 mM sodium phosphate buffer, 20 % glycerol), 15 mM imidazole, 1 % tween 20, 1 mM DTT, 0.1 mM EDTA, and 0.1 mM PMSF at 4 times the cell pellet volume. Suspensions were sonicated on ice for 2 minutes 40 seconds (20 second pulse and 40 second rest cycles). Lysates were centrifuged at 40,000 rpm (164,700 x g) for 20 minutes at speed. Clarified lysates were passed through a 5 mL prepacked Ni-NTA column (GE Life Sciences), washed with 10 column volumes of 15 mM imidazole, then washed further with 5 column volumes of 50 mM imidazole. His_6_-tagged Cyp19 was eluted with 250 mM imidazole. Eluted fractions were concentrated to 0.5 – 1 mL, then injected onto a gel filtration column (Superdex 200 Increase 10/300 GL) with an FPLC AKTA system at 0.7 mL/min in Buffer A. Colored fractions with the highest purity (A10 – A12) were pooled and concentrated (Figure S9).

Cyp19 size was confirmed by a blot analysis with Ni-HRP His probe, and purity was assessed by SDS-PAGE. Briefly, samples were loaded in precast SDS-PAGE gels by standard practices. In blot analyses, protein was transferred for 25 min at 80 V to a FluoroTrans PVDF membrane with the Rapid Transfer (VWR) system. A 3,3’-diaminobenzidine tablet (Sigma) was reconstituted in peroxide/TBS solution for colorimetric detection as recommended by the manufacturers protocol. Sample purity in SDS-PAGE gels were assessed by detection with the Coomassie-based Aquastain (Bulldog Bio).

P450 measurements are detailed by Sohl and Guengerich^50^ with modifications noted here. A stream of nitrogen was used to displace dissolved oxygen in an UltraMicro quartz cuvette (Agilent) prior to adding Na_2_S_2_O_4_ grains (50 μL sample). The cuvette lip was sealed with a parafilm prior to scanning the sample with a single beam UV-Vis spectrometer. A 3-mL Luer-lock syringe with septum was loaded with 1 mL of CO gas, then the septum was replaced with a syringe needle for slowly bubbling CO in the sample solution. The extinction coefficient of 91,000 M^-1^cm^-1^ was used to determine the concentration of P450 from the difference spectrum.

To quantify the binding affinity of Cyp19 for the native substrate androstenedione (ASD), Cyp19 was diluted to 3 μM P450 and titrated with ASD from 1.0 – 46.0 μM. Between each addition of compound, the solution was gently resuspended before absorption scans were measured. An internal reference set at 490 nm corrected for lamp drift during the assay. The peak to trough absorption between 350 and 500 nm from difference spectra were graphed as a function of ASD concentration.
